# Tahoe-x1: Scaling Perturbation-Trained Single-Cell Foundation Models to 3 Billion Parameters

**DOI:** 10.1101/2025.10.23.683759

**Authors:** Shreshth Gandhi, Farnoosh Javadi, Valentine Svensson, Umair Khan, Matthew G. Jones, John Yu, Daniele Merico, Hani Goodarzi, Nima Alidoust

## Abstract

Foundation models have transformed natural language processing and computer vision, yet their potential in single-cell biology—particularly for complex diseases such as cancer— remains underexplored. We present `Tahoe-x1` (Tx1), a family of perturbation-trained single-cell foundation models with up to 3 billion parameters. Tx1 is pretrained on large-scale single-cell transcriptomic datasets, including the *Tahoe-100M* perturbation compendium, and fine-tuned for cancer-relevant tasks. Through architectural optimizations, data loader refinements, and efficient training strategies, Tx1 achieves 3–30× higher compute efficiency than prior implementations of cell-state models. Tx1 jointly learns representations of genes, cells, and compounds using a masked-expression generative objective that incorporates a drug token, enabling flexible adaptation to diverse downstream applications. We evaluate Tx1 across four key disease-relevant benchmarks: (1) prediction of overall and context-specific gene essentiality, (2) identification of genes contributing to the hallmarks of cancer, (3) cell-type classification, and (4) prediction of perturbation responses in held-out cellular contexts. Tx1 achieves state-of-the-art performance across all tasks. We release pretrained checkpoints, training code, and evaluation workflows to accelerate the development of perturbation-trained single-cell foundation models for applications in precision oncology and beyond.

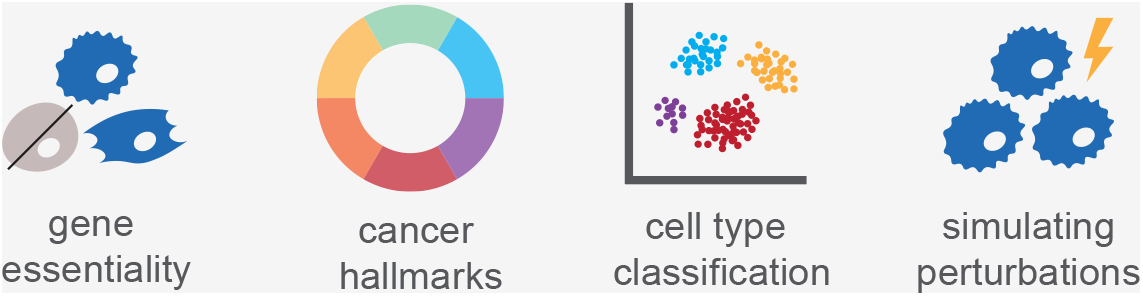

## 1 Introduction

Foundation models, large pretrained architectures capable of transferring to diverse downstream tasks, have transformed natural language processing, computer vision, and other data-rich domains. Their capacity to learn generalizable representations from massive unlabeled corpora and adapt with minimal task-specific supervision has made them indispensable across disciplines.

In biology, recent advances have shown that analogous models trained on large-scale molecular profiles can capture complex biological relationships, enabling applications in functional genomics, cell type annotation, perturbation modeling, and biomarker discovery. The rationale for developing foundation models in cell biology is particularly strong because many experimental settings most relevant to human health, particularly oncology, are inherently data-limited. Clinical samples are difficult to collect, assays are costly, and patient cohorts are often small, often constraining researchers to data-limited regimes. In such contexts, the ability of foundation models to leverage pretraining on a broad set of data and support robust few-shot learning on new tasks and datasets is a decisive advantage.

Single-cell transcriptomics provides an unparalleled view of cellular states and their regulatory programs, offering a high-resolution “language of the cell” from which such models can be learned. The expansion of sequencing throughput, together with initiatives such as the Human Cell Atlas [1], has produced tens of millions of profiles spanning nearly all human tissues, developmental stages, and disease contexts.

Building on these efforts, we at Tahoe Therapeutics recently generated *Tahoe-100M*, the largest single-cell dataset to date, with a focus on perturbing cells from diverse biological contexts [**?** ]. Created using Tahoe’s Mosaic platform, *Tahoe-100M* comprises over 100 million high-quality single-cell transcriptomes from 50 diverse cancer cell lines exposed to more than 1,100 small-molecule perturbations. Unlike most reference atlases, *Tahoe-100M* is explicitly perturbative, capturing cellular responses across a wide range of genetic backgrounds and drug mechanisms of action. This breadth allows models trained on it to better capture causal relationships between genes in varied contexts, moving beyond correlative patterns toward mechanistic insight.

Single-cell foundation models, trained on large corpora of single-cell data such as Tahoe-100M, operate across scales, learning numerical representations for both genes and cells within a shared architecture. This design enables the capture of molecular relationships while also encoding the diversity of cellular states, providing a unified framework for linking genetic features to phenotypes in varied contexts, including cancer.

Here we present `Tahoe-x1` (Tx1), a 3-billion-parameter single-cell foundation model family derived from a scGPT-style encoder architecture [2] and specialized for cancer therapeutics research. Tx1 is trained to jointly model the relationships between genes, pathways, cellular contexts, and chemical perturbations, with particular emphasis on predicting gene essentiality and on producing gene representations that are informative of hallmark pathway membership. These representations reveal that the model has learned biologically meaningful pathway structure, including oncogenic programs, directly from large-scale perturbation data.

Compared with prior single-cell foundation models, Tx1 combines scale, a multi-scale architecture, chemistry-aware conditioning, and perturbation-rich pretraining data with cancer-focused fine-tuning on functional genomics benchmarks such as DepMap essentiality screens and MSigDB hallmark recovery. Key architectural innovations, including transformer training recipes that improve GPU utilization [3, 4], a highly I/O-efficient and streaming capable data-format [5] and efficient distributed training strategies [6] result in our implementation being more than an order of magnitude more efficient than comparable methods, allowing us to train the largest cell-state embedding method to date with comparable computation cost to previous approaches.

Across these evaluations, Tx1 achieves competitive or superior performance relative to smaller models and simple baselines, underscoring the value of scaling model capacity and integrating perturbative data for nuanced functional genomics inference in oncology.

## 2 Results

### 2.1 Architecture, training, and benchmarking of the Tx1 model family

Tx1 models are transformer-based generative architectures trained on single-cell data using a masked gene-expression prediction task that encourages the model to learn gene-gene interactions and context-dependent transcriptional patterns.

Each input cell is represented as a fixed-length sequence of gene tokens, augmented with special tokens denoting the overall cell state (`<cls>`) and, when available, chemical perturbations (`<drug>`). Gene identity, discretized expression level, and masking status are jointly encoded into token embeddings that serve as input to a multi-layer transformer backbone. The model then reconstructs masked expression values through two complementary regression heads: a gene-aware decoder operating on individual token embeddings and a cell-aware decoder that conditions predictions on the global cell embedding (Fig. 1).

**Fig. 1:**
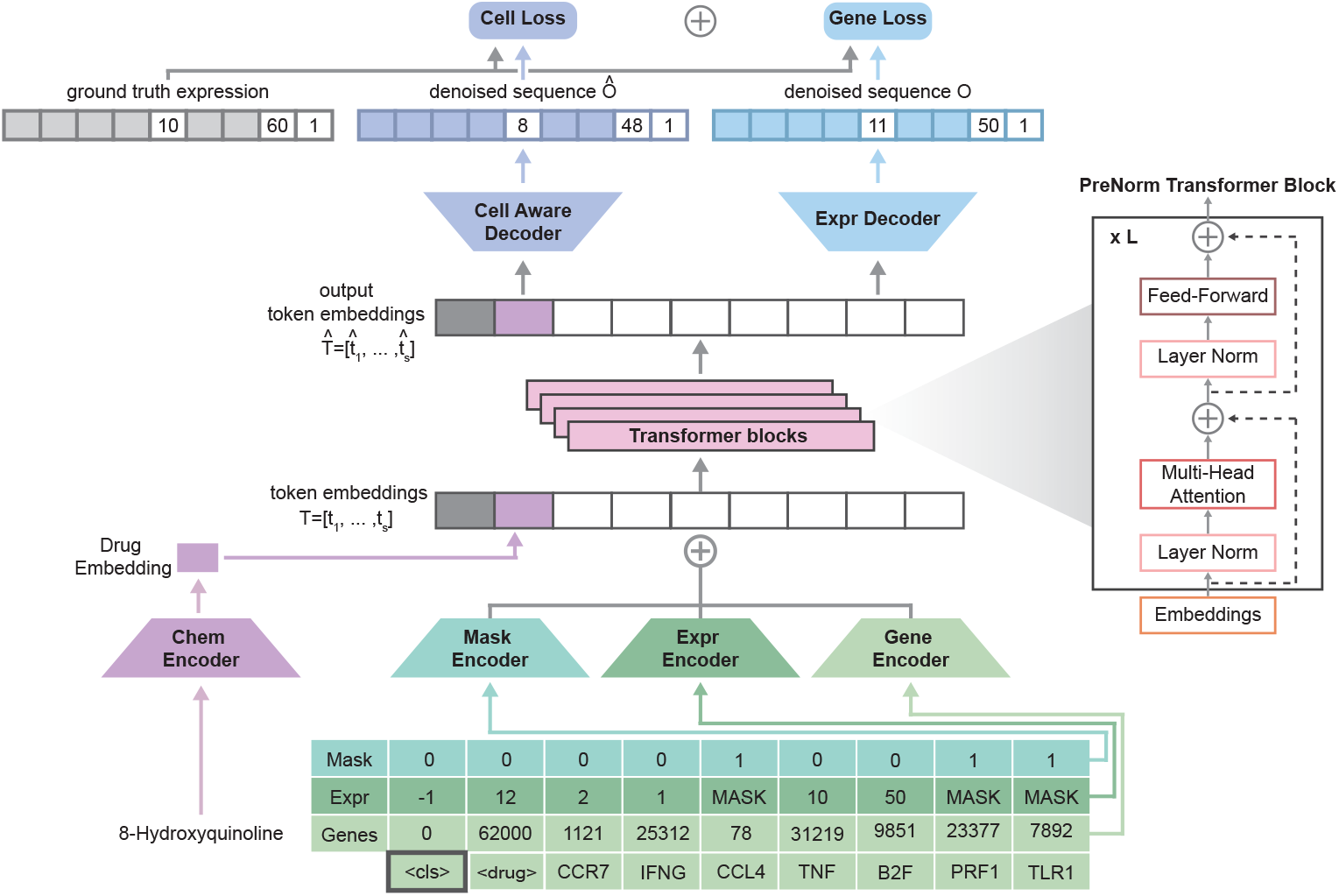
Tx1 model architecture. Schematic of the `Tahoe-x1` (Tx1) model. Each cell is represented as a sequence of gene tokens with associated expression bins and mask indicators. Token embeddings combine information from gene identity, discretized expression level, and a learnable mask vector for masked tokens. An optional `<drug>` token provides chemical context via a chemical encoder initialized with Morgan fingerprints. The resulting embeddings are processed by a stack of prenorm transformer blocks to produce contextualized representations. Two decoders reconstruct masked expression values: the *gene-aware decoder*, which operates on token-level embeddings, and the *cell-aware decoder*, which conditions predictions on the global cell embedding derived from the `<cls>` token. The final objective combines both reconstruction losses equally to form the pretraining objective.

Tx1 was pretrained on up to 266 million single-cell profiles drawn from *Tahoe-100M* [**?** ], Arc scBaseCount [7], and CZ CELLxGENE [8], encompassing both observational and perturbation-rich datasets. The largest model, Tx1-3B, integrates approximately 3 billion parameters across 32 transformer layers and was trained using mixed-precision and fully sharded data parallelism across 128 GPUs.

Training was done in two stages: an initial phase with shorter input sequences (1,024-genes), followed by a second phase of training on a higher quality subset with an increased context length (2,048-genes) sequences to capture higher-order dependencies. Smaller variants (Tx1-70M and Tx1-1.3B) were trained on subsets of the data to assess scaling behavior. Together, these models constitute a unified framework for learning gene, cell, and compound representations from large-scale perturbative single-cell data, enabling generalization across biological contexts and downstream tasks.

We evaluated Tx1 across multiple benchmarks designed to assess its ability to generalize across biological scales and contexts. These include prediction of gene essentiality (both overall and context-specific), recovery of hallmark pathway structure from learned gene representations, cell-type classification across tissues and datasets, and prediction of transcriptional responses to perturbations in previously unseen cellular contexts. Together, these tasks evaluate how well Tx1 embeddings capture the state of a cell, demonstrate their utility for drug discovery, and test the model’s capacity to encode information about gene pathways underlying cancer-related mechanisms.

By enabling prediction of perturbation effects in new contexts where limited or no perturbation data exist, Tx1 advances the goal of context-generalizable cellular modeling. Across benchmarks, Tx1 consistently outperformed prior cell-state embedding methods, highlighting the value of large-scale perturbation pretraining for robust single-cell representation learning.

### 2.2 Representation of transcriptional regulation

#### 2.2.1 Predicting context-specific cancer dependencies using Tx1 embeddings

The prediction of essentiality represents a fundamental challenge in the discovery of cancer targets, with the goal of identifying which genes are required for cancer cell survival in specific contexts. Genes that are essential for specific genotypes or cell states are potential targets for drugging corresponding tumor cells. We used the comprehensive CRISPR-Cas9 screens of the cancer dependency map (DepMap) project in more than 900 cancer cell lines to adapt Tx1 for the prediction of gene essentiality [9]. The model was fine-tuned to predict CERES scores, which quantify the fitness effect of gene knockout while controlling for copy number artifacts [10].

We distinguished between two types of essentiality prediction. *Contextual essentiality* measures context-specific dependencies, such as genes that are essential only in KRAS mutant cells or specific lineages. This task requires the model to learn subtle interactions between genetic background, cellular context, and gene function (Fig. 2A). *Broadly essential genes*, in contrast, are pan-essential genes that are required in all cellular contexts. For both tasks, we used gene expression profiles from the Cancer Cell Line Encyclopedia (CCLE, [11]) as input to obtain gene embeddings from the models and trained a random forest model to predict the corresponding CERES dependency scores. Tx1 demonstrates superior performance in predicting both context-specific and broadly essential genes when compared to other cell state models. It is the only model that performs on the same level as or superior to a linear baseline (Fig. 2B-C). In predicting broadly essential genes, Tx1-3B achieves the strongest mean AUROC and mean AUPRC five times in our evaluation.

**Fig. 2:**
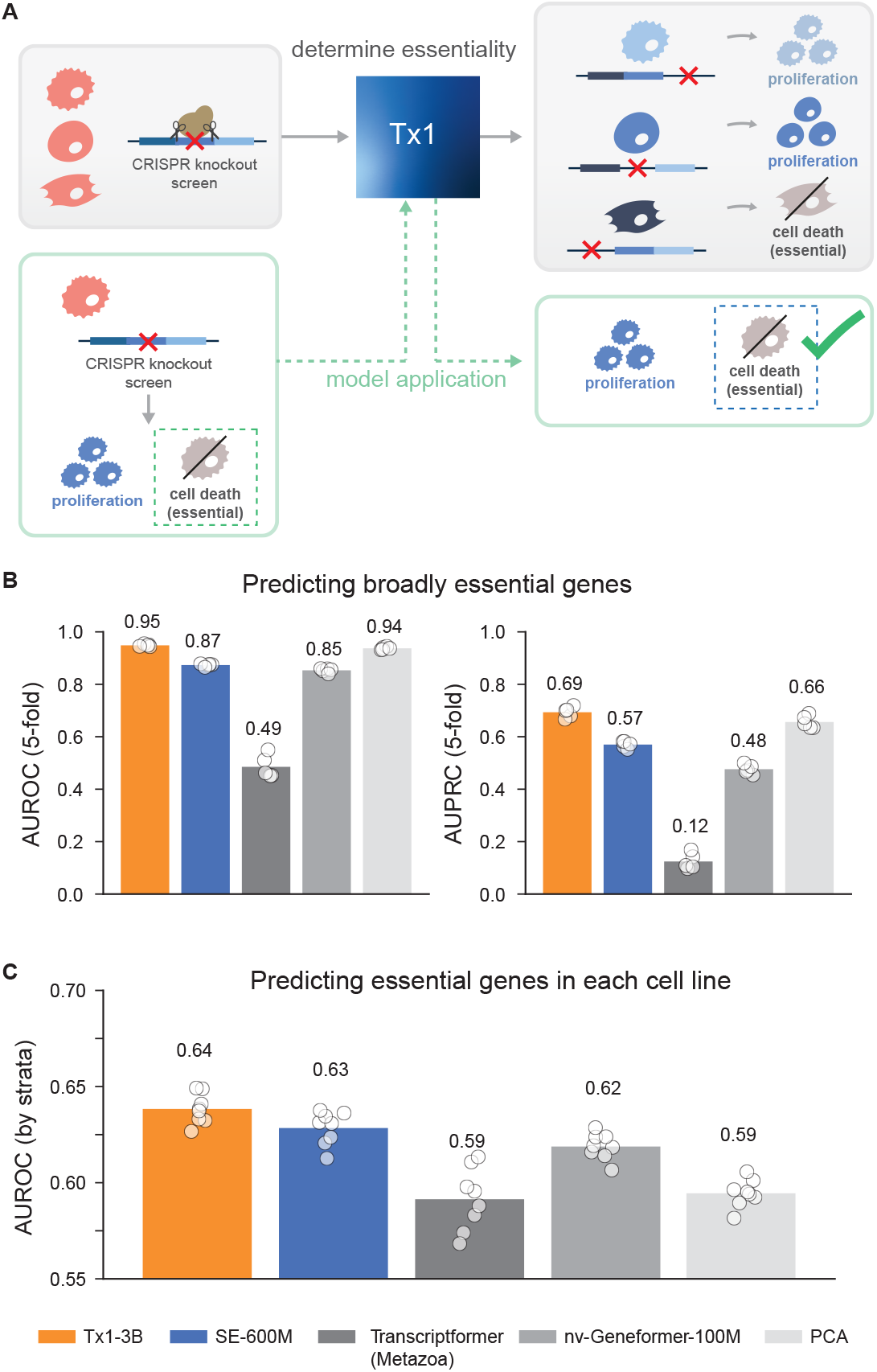
DepMap essentiality benchmarks. (**A**) Schematic of the essentiality prediction framework. Tx1 is fine-tuned on CRISPR knockout screen data from the DepMap project to predict gene essentiality based on transcriptomic profiles. Broadly essential genes are those required for proliferation across most cell lines, whereas context-specific essential genes show selective dependencies in particular cellular or genetic backgrounds. Model validation is performed by comparing predicted essentiality with experimentally measured proliferation phenotypes following gene knockout. (**B**) Performance of Tx1 and baseline models in predicting broadly essential genes, evaluated using five-fold cross-validation. Bars represent mean performance across folds, and individual points show results for each fold. Tx1-3B achieves the highest area under the receiver operating characteristic (AUROC; left) and area under the precision-recall curve (AUPRC; right). (**C**) Context-specific essentiality prediction, evaluated as AUROC stratified by cell line. Tx1-3B achieves the best overall generalization across contexts, indicating its ability to capture cellular dependencies that vary with lineage or genetic background. Together, these results demonstrate that large perturbation-pretrained single-cell foundation models can accurately infer both global and context-specific genetic dependencies.

#### 2.2.2 Predicting hallmark pathway membership using Tx1 embeddings

The MSigDB hallmark collection [12, 13] comprises 50 curated gene sets that capture key biological processes related to cell proliferation and survival, metabolism, immunity, and cancer microenvironment modulation (Fig. 3A). These gene sets are distilled to reduce redundancy typical of other gene set resources, they are widely used in oncology research as they reflect key processes often dysregulated in cancer cells.

**Fig. 3:**
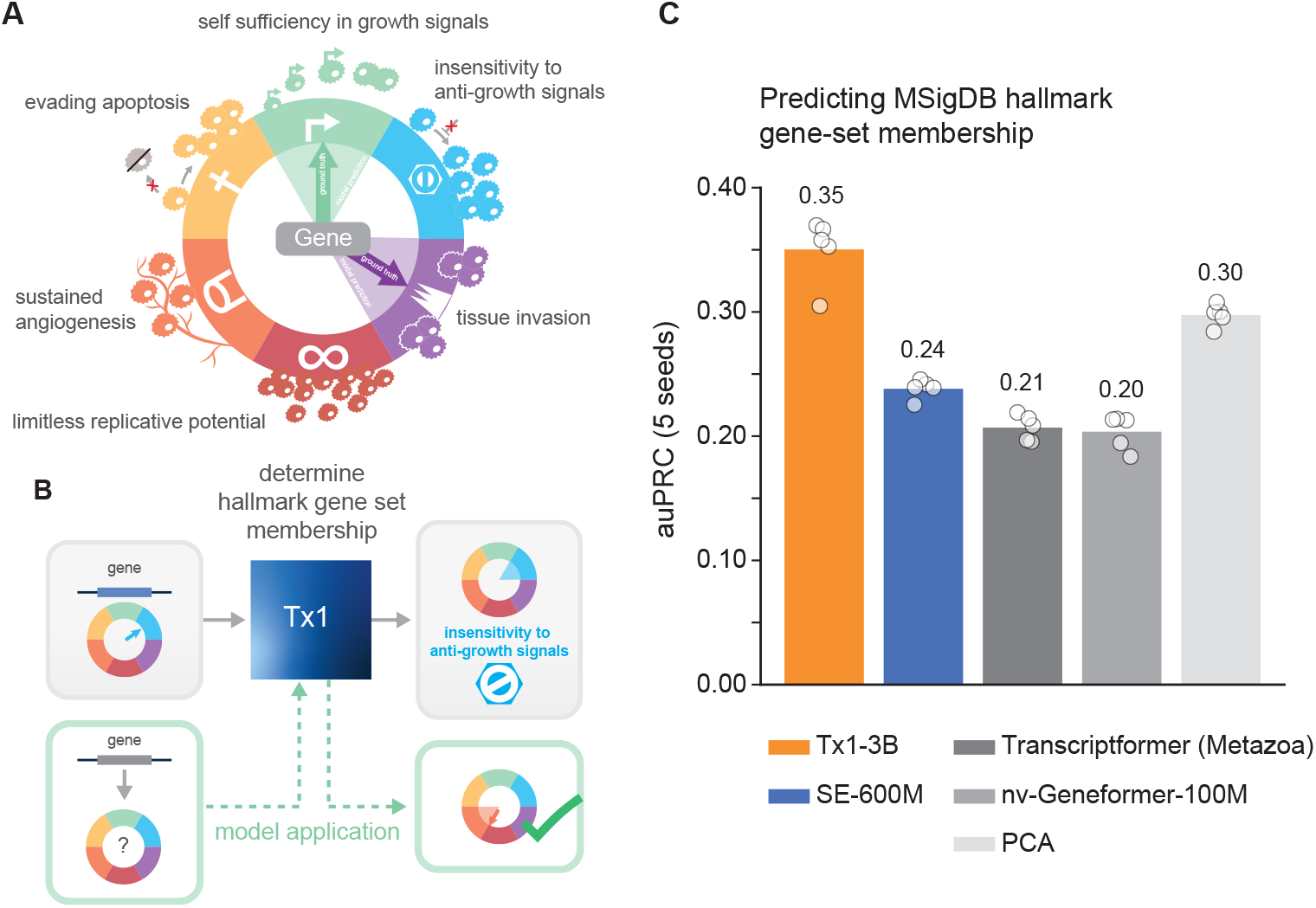
MSigDB hallmark benchmark. (**A**) The canonical *hallmarks of cancer* summarize biological programs that enable tumor growth and survival, including sustained proliferation, evasion of apoptosis, and tissue invasion. The 50 MSigDB hallmark gene-sets capture cancer hallmark-associated as well as other fundamental biological processes. (**B**) Overview of the Tx1 benchmark setup for MSigDB hallmark prediction. Gene embeddings learned by each model are used as input to a lightweight classifier trained to predict a gene’s membership in one or more MSigDB hallmark gene sets. Model performance is validated against curated hallmark annotations. (**C**) Prediction performance (mean area under the precision-recall curve, AUPRC) across five random seeds. Tx1-3B achieves the highest mean AUPRC (0.31), outperforming SE-600M, Transcriptformer (Metazoa), and nv-Geneformer-100M. These results demonstrate that Tx1’s learned gene embeddings capture biologically meaningful structure corresponding to hallmark pathway organization.

We evaluated Tx1’s ability to recover pathway activity from gene expression inputs, testing whether the model’s learned representations capture coordinated gene expression programs. This task is particularly relevant for understanding cancer biology, as many hallmarks represent processes directly related to tumorigenesis and therapeutic response.

Similar to the essentiality benchmark, we use the learned gene embeddings from each of the models and train a shallow MLP to predict a multi-hot label for each gene indicating its membership in one of the hallmark pathways (Fig. 3B). We perform cross-validation and report performance at predicting gene-set membership for held-out genes. Tx1 achieves a mean AUPRC of 0.31 on this task, outperforming other embedding approaches that rely on protein embeddings (Fig. 3C).

### 2.3 Cell state representation

#### 2.3.1 Cell-type classification on Tabula Sapiens 2.0

Although Tx1 was not designed for cell-type classification as a primary objective, we observed that it achieves competitive performance on this benchmark. We obtained cell embeddings on the subset of the Tabula Sapiens dataset [14] used on the public leaderboard and used the *czi-benchmarks* package^1^ to compare the ability of different models to classify cell-type labels provided in the dataset (Fig. 4A). Even though our model was only trained on human cells, we obtain competitive performance with approaches like Transcriptformer that were trained on data from multiple species (Fig. 4B).

**Fig. 4:**
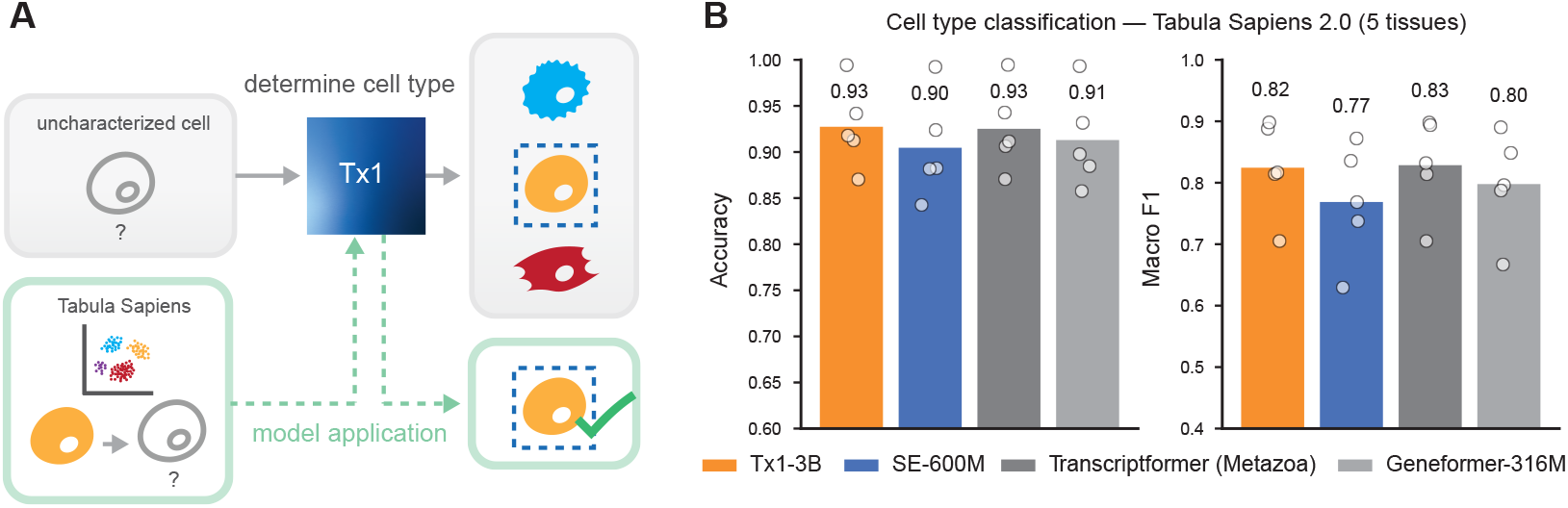
Cell-type classification on Tabula Sapiens 2.0 (CZI benchmark). (**A**) Evaluation of frozen cell embeddings from Tx1 on the Tabula Sapiens 2.0 dataset using the *czi-benchmarks* framework. Embeddings were extracted for cells across five representative tissues (Blood, Bone marrow, Lung, Mammary, and Thymus) and used to train a shallow classifier to predict annotated cell-type labels. (**B**) Comparison of overall performance across models using Accuracy and Macro-F1 metrics. Bars indicate the mean across tissues, while individual dots represent per-tissue results. Tx1-3B achieves performance comparable to or exceeding models trained on multi-species data such as Transcriptformer (Metazoa), despite being trained exclusively on human single-cell data. These results highlight the robustness and cross-tissue generalization of perturbation-pretrained Tx1 embeddings for downstream cell-type classification tasks.

#### 2.3.2 Foundation model embeddings reflect cell state perturbation

A primary utility of cell representation from foundation models is to reflect transcriptional states and in particular subtle changes in the transcriptional state induced by perturbations. Embeddings from foundation models are better able to distinguish treated from untreated cells than typical highly variable gene subset selection (Fig. 5).

**Fig. 5:**
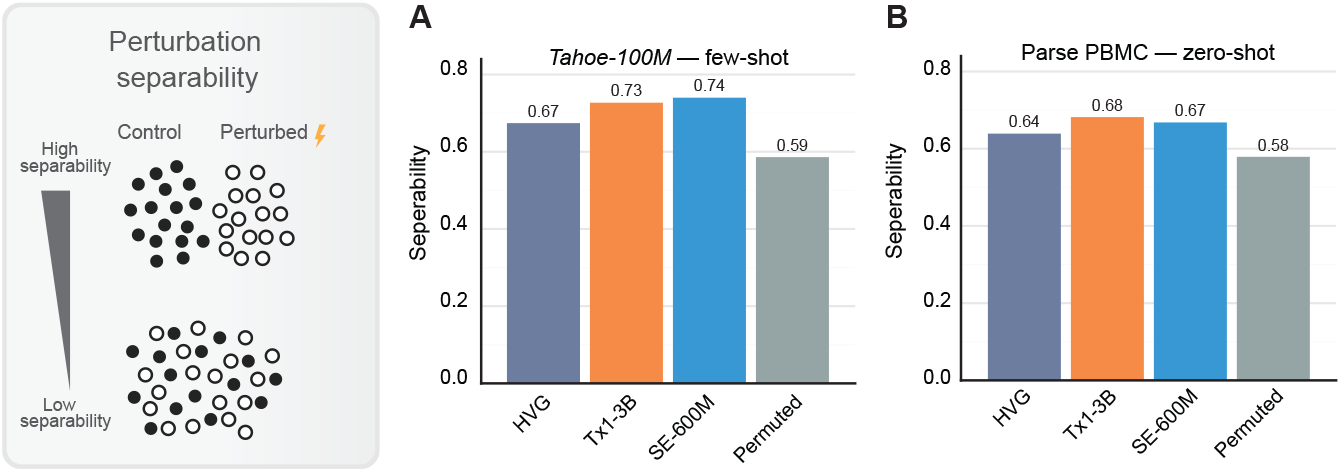
Separability of treated and control cells across representation spaces. Comparison of how well different cell representations capture perturbation-induced transcriptional changes. For each perturbation, separability between treated and untreated (control) cells was quantified using embeddings derived from foundation models versus conventional representations based on highly variable gene (HVG) subsets. Separability was calculated for *Tahoe-100M* (**A**) and *Parse-PBMC* (**B**). Tx1 embeddings exhibit substantially higher separability, indicating that foundation model representations more effectively capture subtle transcriptional shifts associated with cellular perturbations.

### 2.4 Using Tx1 embeddings to predict the effect of perturbations

We demonstrate that Tx1 embeddings can be used as an informative latent space when combined with Arc Institute’s State Transition (ST) model [15] to predict perturbation effects in new cellular contexts (Fig. 6A). Given an unperturbed (control) population, Tx1 maps each cell to an embedding space that organizes transcriptional state while smoothing over assay- and batch-specific noise. We then train the ST module following the hyperparameters reported in the State preprint [15]. The ST module learns how the embeddings shift under a specified perturbation. We quantify performance using Pearson Δ*C* (Methods), which measures correlation between predicted and observed mean expression deltas for a treatment relative to its matched control.

**Fig. 6:**
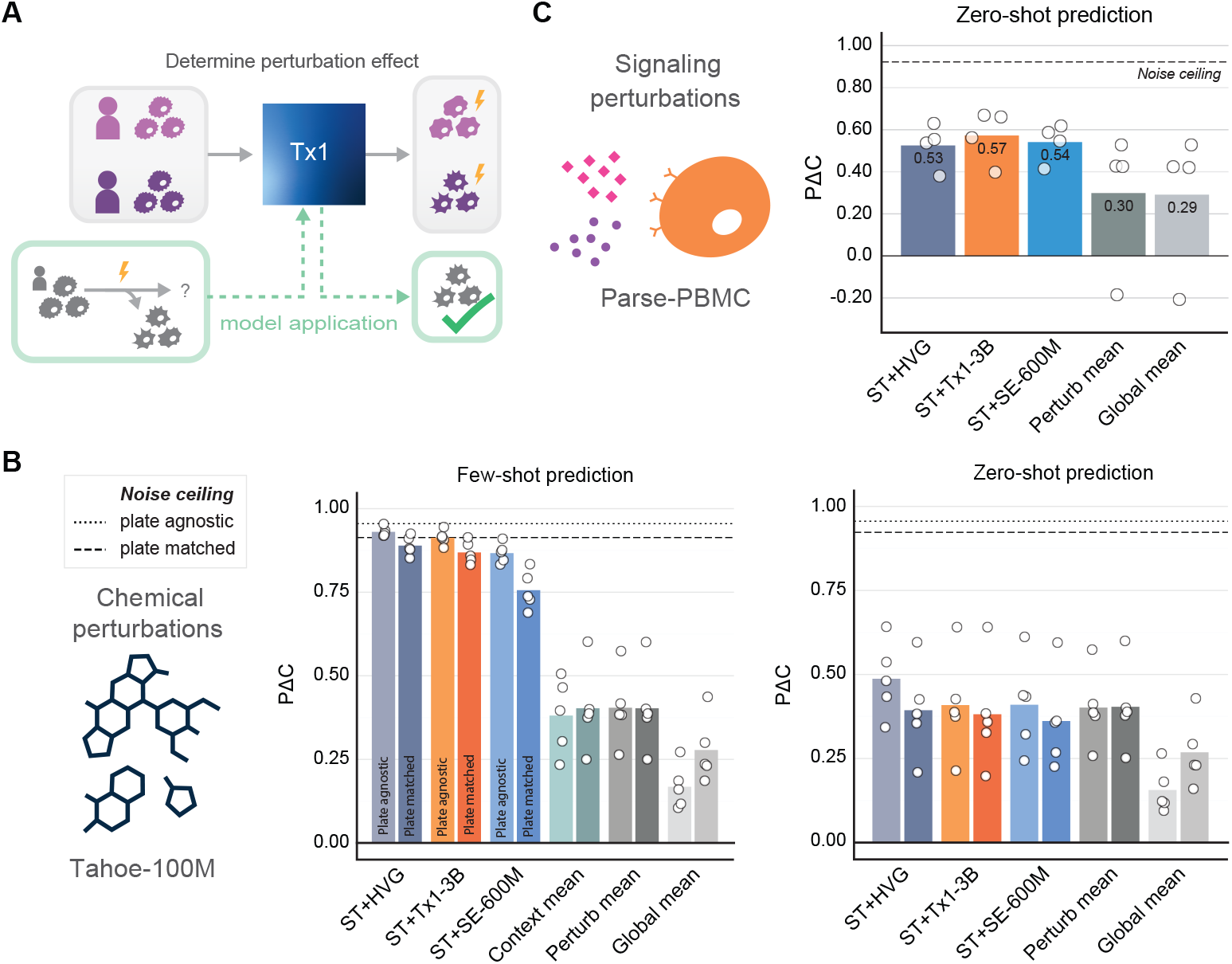
Predicting perturbation effects in held-out cellular contexts using Tx1 embeddings. (**A**) Overview of the Tx1+ST framework for modeling perturbation responses. Tx1 provides smooth, biologically meaningful cell embeddings that capture transcriptional state, which the ST module then learns to transform under specific perturbations. The model is validated by comparing predicted and experimentally observed post-treatment expression profiles. (**B**) Mean perturbation prediction performance on the *Tahoe-100M* dataset in the few-shot prediction task (left) and zero-shot prediction task (right). Pearson ΔC was calculated both without matching treatment and control cells in the same plate (left bars with dashed out-lines) and with plate matching (right bars with solid outlines). Means per cell line context are displayed as white dots. Noise ceiling indicates optimal possible performance in the test set. (**C**) Cross-dataset transfer and adaptation to the Parse PBMC dataset, evaluating zero-shot prediction for held-out donors. Across all settings, Tx1-3B combined with the ST module achieves performance comparable to direct gene-expression modeling and consistently outperforms both SE-600M (from the original State framework) and simpler baselines such as HVG-space, context mean, perturbation mean, and global mean representations. These results demonstrate that Tx1 embeddings provide a robust latent space for learning transferable perturbation-response models across biological contexts.

Two prediction tasks were evaluated on the *Tahoe-100M* data: *few-shot prediction*, where the model sees all contexts and perturbations during training, but not the complete set of combinations, and the more challenging *zero-shot prediction* task, where no data from a subset of cellular contexts is seen during training. Tx1+ST achieves strong performance in the few-shot and zero-shot settings matching the performance of using the gene-expression space directly and surpassing the SE-600M cell-state embedding model developed as part of the STATE model (Fig. 6B).

Generalization to new distributions from initial training data is extremely important for future applications of virtual cell models. To assess performance for this task, we adapted the trained ST models to the *Parse-PBMC* ^2^ dataset in a challenging zero-shot setting that fully leaves out four blood donors. Adapting ST models trained with Tx1 embeddings led to the best performance for predicting how blood from unseen donors react to signaling molecules such as cytokines (Fig. 6C).

### 2.5 Efficient scaling for the Tx1 model series

To understand the relationship between model scale, training efficiency, and downstream performance, we trained the Tx1 model series at three scales: 70M, 1B, and 3B parameters. Fig. 7A shows the training cost versus computational budget (measured in FLOPs) for Tx1 compared to other single-cell foundation models including SE-600M, scGPT, and nv-Geneformer variants. Tx1 achieves substantially improved training efficiency, with 3–30× better compute efficiency relative to these prior models. This improvement stems from several architectural and implementation choices, most notably the removal of custom attention masking that enables efficient kernels such as FlashAttention, mixed-precision training, and optimized distributed training strategies using Fully Sharded Data Parallel (FSDP).

**Fig. 7:**
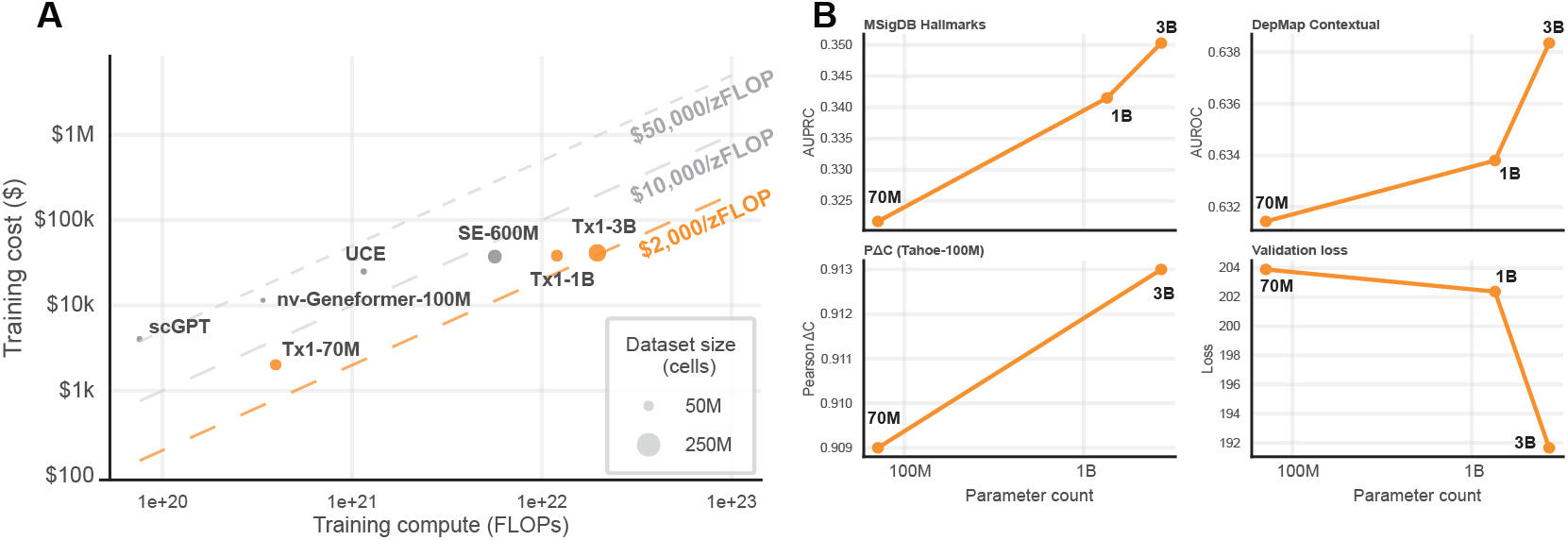
Compute-efficient scaling and performance gains in the Tx1 model series. (**A**) Training cost versus computational budget (measured in FLOPs) for the Tx1 models compared with prior single-cell foundation models, including SE-600M, UCE, scGPT, and nv-Geneformer-100M. Point size corresponds to the number of single cells used for pretraining. Tx1 achieves 3–30× higher compute-efficiency relative to these models, largely due to optimized implementation choices such as the removal of custom attention masking, adoption of FlashAttention kernels, mixed-precision training, and distributed training using Fully Sharded Data Parallel (FSDP). (**B**) Scaling trends for downstream performance across tasks and model sizes (70M, 1B, and 3B parameters). Increasing scale consistently improves results across diverse benchmarks: MSigDB Hallmark pathway prediction (AUPRC), DepMap contextual essentiality prediction (AUROC), and perturbation effect modeling on *Tahoe-100M* (Pearson Δ*C*). Validation loss similarly decreases with model scale, confirming that larger models capture richer structure in the pretraining data. Together, these results demonstrate that efficient training enables practical scaling of multi-billion-parameter single-cell foundation models while maintaining favorable cost-performance tradeoffs.

The efficiency gains are particularly important for enabling the scaling to 3B parameters, which would otherwise be prohibitively expensive. By reducing training costs, we can explore larger model capacities and longer training runs that translate into improved downstream performance. Fig. 7B demonstrates that increasing model scale yields consistent improvements across multiple benchmarks. On MSigDB Hallmarks and DepMap contextual essentiality prediction, performance increases monotonically with parameter count. For the *Tahoe-100M* state transition task (PΔC), we observe substantial gains from 70M to 3B parameters. Validation loss also decreases consistently with scale, confirming that larger models better capture the underlying transcriptomic patterns in the pretraining data.

These results demonstrate that efficient training infrastructure, combined with thoughtful architectural choices, enables the practical development of multi-billion parameter single-cell foundation models. The sustained performance improvements across scales suggest that further scaling beyond 3B parameters may yield additional gains, particularly for complex context-dependent prediction tasks in precision oncology, and as large single cell atlases of 1B cells and beyond become available.

## 3 Discussion

Tx1 demonstrates that scaling single-cell foundation models, particularly when trained on large perturbative datasets such as *Tahoe-100M*, can deliver tangible improvements in cancer-relevant functional genomics tasks. By learning numerical representations for both genes and cells within a unified architecture, the model integrates molecular and cellular scales of information, enabling it to capture context-dependent genetic dependencies and encode biologically meaningful pathway structure. Tx1’s gene embeddings being predictive of hallmark pathway membership suggests that the model internalizes higher-order biological programs directly from transcriptomic perturbation data, without the need for explicit pathway supervision.

A critical insight from the Arc Institute’s State model is that learning information-rich cell embeddings is essential for context generalization, defined as the ability to make accurate predictions in cellular contexts not seen during training. This capability is particularly important for translational applications in oncology, where functional data such as drug sensitivity can often only be collected in simplified or incomplete models, such as cell lines *in vitro* or patient-derived xenografts lacking an intact immune system. If a model can generalize across contexts, it becomes possible to transfer knowledge from these tractable systems to make reliable predictions of how a tumor in a patient will respond, given deep molecular profiling of a biopsy from that tumor. Tx1’s improvements in marginal essentiality prediction, along with its strong performance in recovering pathway structure, suggest that it is beginning to achieve this kind of generalization, enabling more effective transfer learning from experimental models to patient settings.

*Tahoe-100M* played a central role in enabling this capability. Earlier iterations of these architectures, trained solely on large observational compendia, did not achieve similar performance on context-dependent prediction tasks. By contrast, pretraining on the large, diverse, and explicitly perturbative *Tahoe-100M* dataset allowed Tx1 to observe a far broader range of context–perturbation combinations and learn causal relationships that generalize beyond specific drugs, targets, or cell types. This demonstrates that large-scale perturbation data is not simply helpful but essential for pushing foundation models toward causal and translational utility. *Tahoe-100M* is, by a wide margin, the largest single-cell dataset generated to date, and by releasing it publicly we aimed to catalyze similar efforts across the community. Progress in this space will depend on sustained expansion of perturbative datasets that match or exceed this scale, covering new cell types, perturbation modalities, and disease contexts.

More broadly, Tx1 illustrates the potential for single-cell foundation models to operate effectively in data-scarce experimental regimes. Because many translational applications involve small patient cohorts or rare subtypes, the ability to adapt to new contexts with few-shot learning is critical. Pretraining on massive, diverse datasets such as *Tahoe-100M* provides a foundation that can be fine-tuned or prompted to handle novel perturbations, genetic backgrounds, or phenotypic readouts with minimal additional data.

Practically, this enables two oncology workflows: (i) *screen-to-patient transfer*, using Tx1+ST trained on cell-line screens to prioritize compounds in tumor contexts lacking perturbation data; and (ii) *context expansion*, extrapolating effects from a small set of sentinel lines to related, unperturbed lines. In both cases, Tx1’s smoother latent geometry pairs naturally with the transport model, yielding more accurate effect-size ranking in new cellular contexts.

There are, however, limitations to the current approach. While *Tahoe-100M* is broad and deep within the domain of *in vitro* cancer cell line models, it does not yet encompass the full range of primary cell types, tissue microenvironments, or *in vivo* perturbations that will be necessary to capture the complexity of patient tumors. Similarly, while the current architecture integrates gene and cell representations, it does not yet incorporate other molecular modalities such as protein abundance or spatial context, which could further enrich its biological understanding. Beyond scaling the model size, expanding both the diversity of pretraining data and the range of integrated modalities represents a clear next step.

Ultimately, `Tahoe-x1` points toward a future in which large, multi-scale, perturbation-trained single-cell foundation models become core infrastructure for systems biology and therapeutic discovery. As the scale and diversity of available perturbative data continue to grow, and as modeling architectures evolve to incorporate richer biological priors, these models will be positioned to bridge the gap between cellular-scale measurements and actionable insights for precision medicine.

## 4 Methods

Tahoe-x1 (Tx1) is a transformer-based foundation model trained on single-cell data via masked gene-expression prediction. The task encourages the model to learn gene-gene interactions and correlations within each sequence. Fig. 1 summarizes the architecture as follows:

1. We form a sequence consisting of special tokens and sampled gene tokens;
2. Raw counts are discretized into B expression bins;
3. Token embeddings are obtained by summing a gene-identity embedding, an expression embedding, and (for masked positions) a learnable mask embedding;
4. When available, a chemical token provides drug context;
5. A stack of transformer blocks produces contextualized gene representations;
6. Lastly, two regression heads decode masked expression values—one directly from gene-aware token embeddings (gene-aware) and one from the cell embedding alone (cell-aware). Training uses a masked denoising objective combining both heads with equal weight.

In the sections that follow, we discuss each component in detail.

### 4.1 Modules

#### 4.1.1 Sequence Formation

As a first step, we construct, for each cell, a sequence of genes with fixed length S. We randomly sample genes from the set of expressed genes and discretize their expression values into B bins (Section 4.1.2). If the number of expressed genes is smaller than the target number of gene tokens, we pad the sequence with the special token `<pad_token>`. We also include two special tokens: `<cls>` to capture the overall cell state, and `<drug>` to encode drug information when such metadata are available (e.g., in *Tahoe-100M* ). The expression values associated with these special tokens are initialized to a fixed unused scalar. For convenience, we standardize the order so that the first token is `<cls>`, the second token is `<drug>` (if present), followed by one token per selected gene; any remaining positions are filled with `<pad_token>`. We denote the final sequence length by S.

#### 4.1.2 Binning

We follow a similar expression binning strategy as was proposed in scGPT [2]. The procedure is as follows:

1. Define a vector of equal spaced B grades Z = [0, …, 1] where B corresponds to the number of bins.
2. For a vector of raw counts 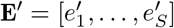, compute bins b as the quantiles of **E*′*** for each grade *Z*. Thus, b is the 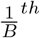 quantile of **E*′*** and b is the maxima of **E*′***. To compute the quantile, we map each grade in **Z** to the range of indices in **E*′*** to find the location of the quantile in the sorted input. If the quantile lies between elements in **E*′***, we use linear interpolation based on the fractional part of the quantile index.
3. Compute vector of digitized expression values 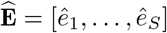, with each value corresponding to the index of the bin in which it belongs, according to one of the following rules:

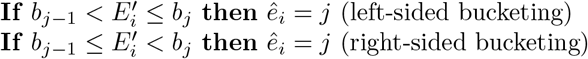

Left-sided bucketing returns the **first** index *j* that satisfies the criteria above, whereas right-sided returns the last index that meets the criteria. If no suitable index is found (such as for the smallest element when using left-sided bucketing) then an index of 0 is returned.

We opted for the left-sided binning approach and add a constant value of 1 to the binned counts so that the dynamic range for both left and right-sided binning is the same, ie 1, …, **B** − 1. This differs from the scGPT paper, which used right-sided binning by default. The consequence of right-sided binning is that small counts get inflated to higher bins and the overall dynamic range of the binned values is smaller.

#### 4.1.3 Masking

From the constructed sequence, we randomly mask a fraction of tokens according to a masking ratio **P**. We define a binary mask vector :

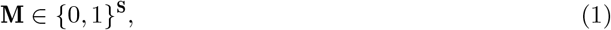

where *M*_*i*_ = 1 if the expression value at position *i* is hidden from the model, and *M*_*i*_ = 0 otherwise. Special tokens `<cls>` and `<drug>` are never masked. For masked positions, the corresponding digitized expression values are replaced by a sentinel value (e.g., −1), preventing the model from accessing the true expression and cheating during training. We experimented with masking ratios 15%, 30%, 50%, 75% and found that 30% and 50% yielded the best performance. All Tx1 models were trained with a masking ratio of 50%.

#### 4.1.4 Gene Encoder

The model maintains a vocabulary that maps Ensembl gene identifiers to unique integer indices. The Tx1 vocabulary includes approximately 67,000 genes. Unlike related methods such as STATE [15], which restrict the model to protein-coding genes only, our vocabulary spans both protein-coding and non-coding genes. For each sequence of length S, the gene tokens are represented by their vocabulary indices

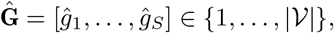

where |𝒱| denotes the vocabulary size. These indices 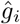 are mapped to continuous embeddings by the Gene Encoder ℱ_GeneEnc_, which consists of a randomly initialized embedding layer ∈ ℝ^|𝒱|*×d*^ followed by a LayerNorm operation:

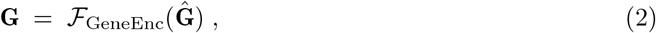

where **G** = [*g*_1_, *…,g*_s_] where G = [g1,∈ ℝ^**S**×*d*^,the resulting *d*-dimensional gene identity embeddings

We also experimented with augmenting these learnable embeddings using external representations such as ESM2 [16] embeddings for protein-coding genes. However, the final Tx1 series employs only the learnable embedding layer, as this configuration yielded simpler training and competitive performance.

#### 4.1.5 Expression Encoder

Digitized expression values 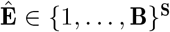 (or the mask sentinel at masked positions) are mapped to *d*-dimensional embeddings via ℱ_ExprEnc_, which is a small MLP with nonlinear activation function and dropout (rate 0.1), followed by LayerNorm.

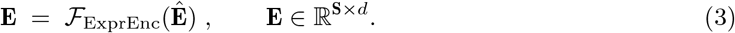

#### 4.1.6 Mask Encoding

To explicitly mark masked positions, we learn a vector 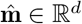 named as a mask vector and broadcast it to masked tokens yielding 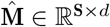 as below:

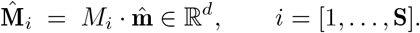

Implied by formula, 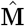 masked tokens. would simply be zero on unmasked tokens and equal to the vector 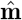 on

##### Token embeddings

The final token embeddings **T** = [*t*_1_, …, *t*_S_] that is input to the transformer is sum of the three components:

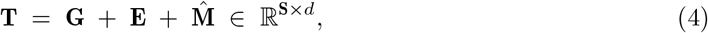

which compactly encodes gene identity, discretized expression, and mask status.

#### 4.1.7 Chemical Encoder

The Tx1 series optionally incorporates drug information alongside with gene identity and expression values. The chemical information is injected into the model via an extra token `⟨drug⟩` to help the model learn a richer cell and gene embeddings.

To this purpose, we introduced ℱ_ChemEnc_, which consists of an embedding layer and a MLP layer with nonlinear activation function and a LayerNorm on top. We initialize the drug-embedding layer with 2048-dimensional Morgan fingerprints for all drugs available in the training corpus (Tahoe100M is the only dataset in our training set that includes drug metadata, comprising 377 drugs.)

First, we create a one-hot vector of size equal to the total number of drugs, 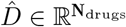, based on the drug index. We reserve index 0 and initialize it with an all-zero vector for cells that do not include chemical information. The vector 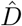 serves as the input to the chemical encoder. Within the chemical encoder, the one-hot encoding of a drug is first mapped to its fingerprint representation through an embedding layer, producing a 2048-dimensional vector. The index 0 is ignored by the embedding layer for samples without drug information, resulting in a 2048-dimensional all-zero vector. The fingerprint representation is then projected to dimension *d* by ℱ_ChemEnc_. The resulting *d*-dimensional embedding captures the chemical information and is injected into the sequence to enable richer joint representations of cells and genes. By design, the second token position in the sequence is optionally reserved for the `<drug>` token.

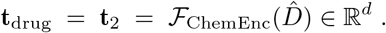

In addition, the chemical embedding layer is fully trainable and not frozen, allowing drug representations to be refined throughout pretraining.

#### 4.1.8 Transformer

Once we have the total token embeddings **T** (including the drug embedding) we pass them to *L* layers of a standard Pre-Norm transformer [3]. The transformer layers ℱ_Transf_ learn token-to-token relationships and produce the final contextualized embeddings 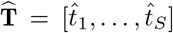 which compactly capture gene-gene interactions and higher-order dependencies.

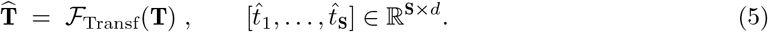

We adopted the Pre-Norm variant of transformer block as shown in Fig.1 for improved gradient flow and training stability compared to Post-Norm [17]. We also used expansion ratio of 4 for FFN layers and LayerNorm as the normalization layer.

#### 4.1.9 Expression Decoder

Given the contextualized token embeddings 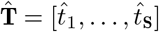 from the transformer, we predict digitized expression values using the Expression Decoder ℱ_ExprDec_. This module consists of a one-layer MLP with hidden size *d*, nonlinear activation, and output dimension 1, followed by LayerNorm.

Formally,

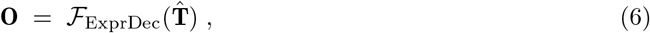

where **O** ∈ ℝ^**S**^ represent the predicted expression values.

##### Gene Loss

Training uses a masked regression objective computed only over masked positions within the sequence. Let 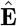 be the ground-truth binned expressions and **M** ∈ {0, 1}^**S**^ the mask vector. The gene loss is the mean squared error (MSE) of 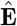 and **O** on masked locations as follow:

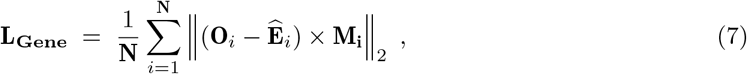

where N is the total number of cells within the dataset.

We refer to this as *gene loss* because, when predicting expression values, the model conditions on the gene tokens of the **entire sequence**, including the `<cls>` and `<drug>` tokens, as contextual information. This formulation penalizes prediction errors only on masked tokens. We also experimented with classification into B discrete bins using cross-entropy, but MSE regression performed better, likely because bin indices have an inherent ordering that cross-entropy does not exploit and treats them as an independent classes.

#### 4.1.10 Cell Aware Decoder

In addition to token-wise prediction, the model also includes a cell-aware decoder that uses only the cell embedding to reconstruct masked expressions by implementing a bilinear attention mechanism. The cell embedding is defined as the contextualized output of the ℱ_Transf_ for `<cls>` token:

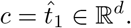

Given *c* and token embeddings 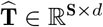, the decoder computes query vectors as:

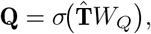

where *W*_*Q*_ ∈ ℝ^*d×d*^ is a learned projection and *σ* denotes a sigmoid activation.

These query vectors are then projected through a weight matrix *W*_*O*_ ∈ ℝ^*d×d*^ and combined with the cell embedding *c* via matrix multiplication to produce expression predictions for all genes simultaneously,

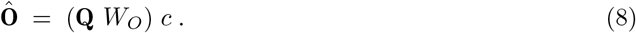

This bilinear form enables the model to generate cell-specific gene expression predictions by modulating gene representations through learned query transformations that are then matched against the global cell context. The resulting inner product operation produces a compatibility score between each gene’s query representation and the cell’s embedding, effectively modeling how each gene’s expression should be influenced by the overall cellular state captured in the cell embedding.

##### Cell Loss

We introduce *cell loss* to jointly train the Cell-Aware Decoder with the rest of the network and to encourage the cell embedding to capture the global state of the sequence. Similar to gene loss, cell loss is computed only on masked tokens. We refer to it as cell loss because, for each query gene, the model relies exclusively on the cell context *c*. Formally, the loss is defined as the mean squared error (MSE) over masked positions:

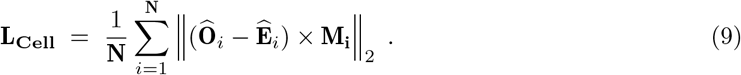

As with gene loss, we use regression with MSE rather than classification with cross-entropy, since the discretized bins have an inherent ordering that is not captured by categorical loss.

#### 4.1.11 Pretraining Objective

The final training objective combines the cell loss and gene loss described in Sections 4.1.10 and 4.1.9.

This formulation is analogous to sequence denoising objectives used in masked language modeling and models such as BERT[18] and RoBERTa[19].

Since both losses are MSE-based denoising objectives applied to masked tokens, and their scales are numerically comparable, we assign them equal weight so that each contributes equally to training. The total loss is defined as:

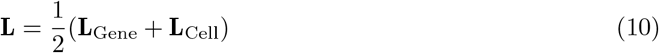

##### Summary

Tab.1 summarizes all the sub-modules within the Tx1 models and their architecture and input and output dimensions.

**Table 1:**
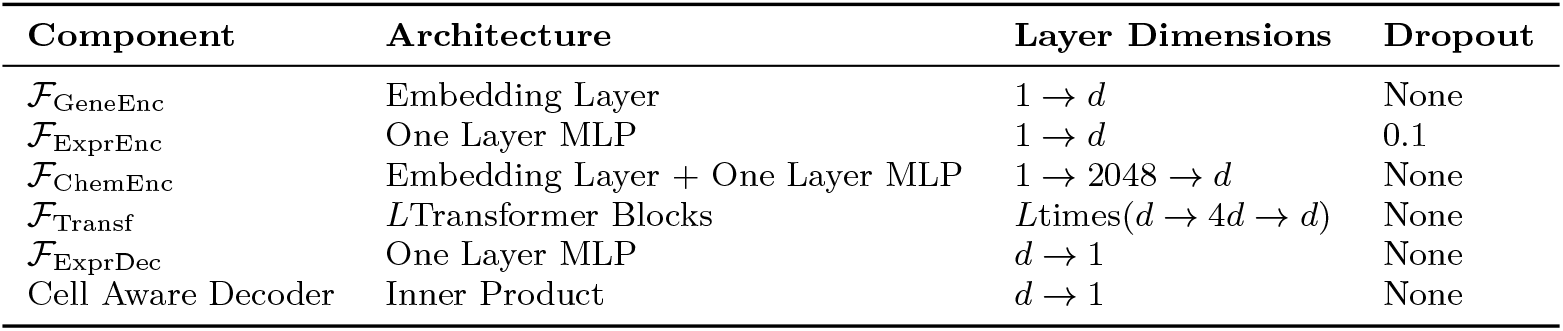
Architectural details for TX modules.

| Component | Architecture | Layer Dimensions | Dropout |
| --- | --- | --- | --- |
| $\mathcal{F}_{\text{GeneEnc}}$ | Embedding Layer | $1 \rightarrow d$ | None |
| $\mathcal{F}_{\text{ExprEnc}}$ | One Layer MLP | $1 \rightarrow d$ | 0.1 |
| $\mathcal{F}_{\text{ChemEnc}}$ | Embedding Layer + One Layer MLP | $1 \rightarrow 2048 \rightarrow d$ | None |
| $\mathcal{F}_{\text{Transf}}$ | $L$Transformer Blocks | $L \text{times}(d \rightarrow 4d \rightarrow d)$ | None |
| $\mathcal{F}_{\text{ExprDec}}$ | One Layer MLP | $d \rightarrow 1$ | None |
| Cell Aware Decoder | Inner Product | $d \rightarrow 1$ | None |

### 4.2 Pretraining

#### 4.2.1 Datasets

We used three datasets for pretraining the Tx1 series: the Tahoe-100M dataset [20], Arc scBaseCount [21], and CZ CELLxGENE [22]. Each dataset was split into a training set, used to fit the model; and a validation set, used to compute validation loss and evaluation metrics. The number of cells drawn from each dataset for pretraining is summarized in Tab. 2.

We trained our largest model, Tx1-3B, on all three datasets, while the smaller versions were trained before scBaseCount was released, and thus were trained only on the Tahoe-100M and CELLxGENE datasets.

For the Tahoe-100M dataset, we included all cells that passed the ‘full’ quality filters and retained all genes expressed in at least one cell. For CELLxGENE data, we used the January 21, 2025 release, restricted to primary datasets to eliminate duplicates, and included genes with a count of at least 10 per 10,000-cell chunk. For scBaseCount, we included cells with at least 10 expressed genes and 20 total counts. In addition, we created a filtered version of each dataset with more stringent thresholds requiring at least 1,000 expressed genes and 2,000 total counts per cell which we used for the final refinement epoch of Tx1-3B training.

#### 4.2.2 Pretraining Recipe

To simplify multi-node distributed training, Tx1 series are implemented in the Composer library^3^, which is built on top of PyTorch^4^. To make pretraining efficient, we employed mixed-precision training with BF16 data type [23], which stores some parameters and intermediate values in full precision while others are stored in half precision, thereby reducing memory usage. We also used Fully Sharded Data Parallel (FSDP) with the full-shard strategy [6], which partitions data, model weights, and optimizer states across multiple GPUs. This enabled us to effectively train the largest model (Tx1-3B) across 128 GPUs.

All model scales use a shared embedding size *d*, which is consistent across encoders, transformer blocks, and decoder inputs. Following common practice in large language models [17, 18, 24], we used an MLP expansion ratio of 4 within the transformer block. We also adopted a lower-precision LayerNorm implementation for faster computation. Gradient accumulation was applied, which allowed us to train with very large batch sizes without running out of memory.

We used AdamW [25] as the optimizer, weight decay for regularization, cosine annealing as the learning-rate scheduler, and a 5% linear warmup schedule.

Tx1-70M and Tx1-1.3B were trained without the chemical token `<drug>`, while Tx1-3B was trained with the chemical token injected into the sequence, allowing the model to integrate drug information as contextual input for cell and gene embeddings.

We trained the models on GPUs provisioned across multiple platforms, including Databricks MosaicML, Google Cloud, Oracle Cloud Infrastructure (OCI), and Lambda Labs.

##### Attention Mask

We improved the attention masking implementation of scGPT [2] noticeably in two steps. In scGPT model, the attention computation involved two sequential attention calls, as shown in Fig. 8A: one to the Torch backend for cross-attention (slow) and another to the FlashAttention backend for self-attention (faster). These two separate sequential calls were inefficient and slow.

**Fig. 8:**
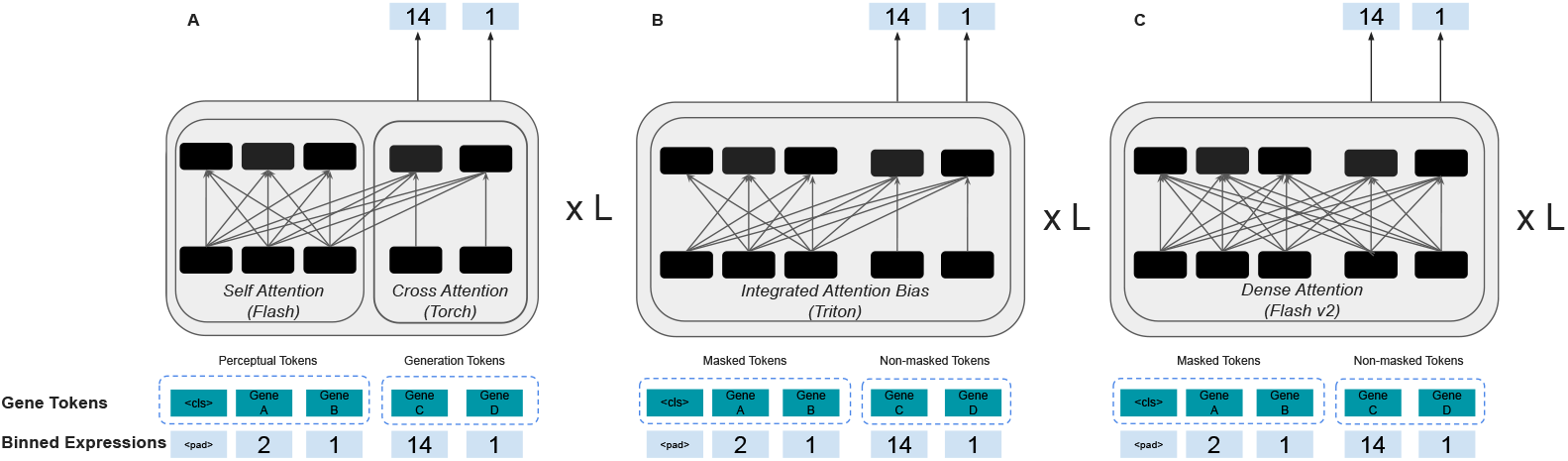
Attention masking strategies. (**A**) The original masking strategy in scGPT, which required two separate attention calls: one for self-attention using FlashAttention and another for cross-attention using the Torch backend. (**B**) The unified masking implementation that merges these two attention passes into a single masked attention call using the Triton backend. (**C**) The final attention implementation used for training Tx1-3B, employing FlashAttention with dense attention across all tokens, allowing full token-to-token interactions without custom masking.

In the first step of optimization, we merged both attention passes into a single, consolidated masked attention call as shown in Fig. 8B. This was achieved by introducing a bias matrix directly into the attention weights that encoded the masking information. Implemented through the Triton backend, this modification substantially increased throughput and cut GPU memory use during the forward pass by about 10× while still enforcing masking. This design was used for efficiently training Tx1-70M and Tx1-1B.

In the second step, we pushed the idea further and investigated whether the custom attention mask was necessary for performance. The attention mask strategy in scGPT only allowed masked (generation) tokens to attend to themselves and to unmasked (perceptual) tokens. However, this appeared redundant, as the expression values of masked tokens were already padded and hidden from the model. Consequently, the masking provided no additional benefit, and theoretically, all tokens could attend to each other without data leakage. This insight enabled us to remove the custom mask entirely and apply dense gene-to-gene attention as shown in Fig. 8C. Since FlashAttention[26, 27] backend does not support custom masks, this change allowed us to use FlashAttention for the entire sequence, further improving throughput and training speed without any degradation in model performance.

The Tx1-70M and Tx1-1.3B models were trained using the Triton backend as shown in Fig. 8B. For Tx1-3B, we removed the custom attention mask and adopted dense attention across all tokens, as shown in Fig. 8C. This modification allowed both masked and unmasked tokens to attend to every token in the sequence, fully eliminating the custom mask used in scGPT and enabling the use of FlashAttention throughout.

##### Continued Training

Following multi-stage training strategies used in LLMs [28, 29], Tx1-3B was trained in two stages. In the first stage, the model was trained with a sequence length of 1, 024 for four epochs on the datasets described in Tab. 2. In the second stage, training continued with an increased sequence length of 2, 048 for one additional epoch on the filtered datasets. Unlike the first stage, where we used a learning rate (LR) of 3 × 10^*−*4^ with a cosine annealing scheduler and warmup, the second stage used a constant learning rate of 3 × 10^*−*5^ (equal to the LR at the end of the first stage). As described in (Section 4.2.1) we apply more stringent quality filters to the data for Stage-2 training to ensure we remove cells with too few genes to take advantage of the increased context length.

**Table 2:** Total number of cells for datasets used in pretraining of Tx1 series.

| Dataset | # Training Cells | # Validation Cells |
| --- | --- | --- |
| Tahoe-100M [20] | 94,668,090 | 956,244 |
| Arc scBaseCount [21] | 111,220,664 | 921,578 |
| CZ CELLxGENE [22] | 60,746,795 | 613,604 |
| **Total** | **266,635,549** | **2,491,426** |

##### Hyperparameters

Details of our hyperparameter choices for each model are provided in Tab. 3.

**Table 3:**
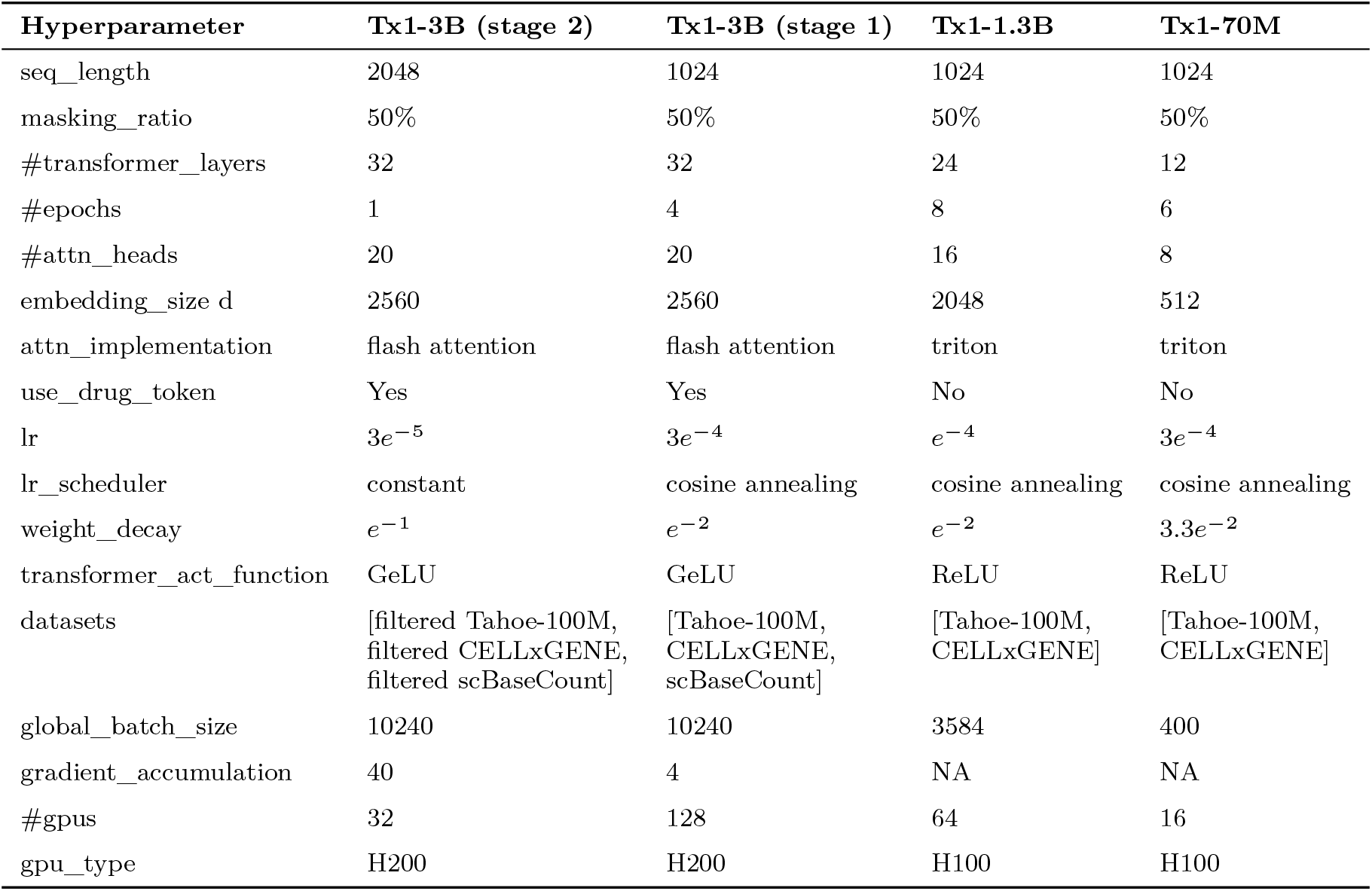
Hyperparameter choices per model.

| Hyperparameter | Tx1-3B (stage 2) | Tx1-3B (stage 1) | Tx1-1.3B | Tx1-70M |
| --- | --- | --- | --- | --- |
| seq_length | 2048 | 1024 | 1024 | 1024 |
| masking_ratio | 50% | 50% | 50% | 50% |
| #transformer_layers | 32 | 32 | 24 | 12 |
| #epochs | 1 | 4 | 8 | 6 |
| #attn_heads | 20 | 20 | 16 | 8 |
| embedding_size d | 2560 | 2560 | 2048 | 512 |
| attn_implementation | flash attention | flash attention | triton | triton |
| use_drug_token | Yes | Yes | No | No |
| lr | $3e^{-5}$ | $3e^{-4}$ | $e^{-4}$ | $3e^{-4}$ |
| lr_scheduler | constant | cosine annealing | cosine annealing | cosine annealing |
| weight_decay | $e^{-1}$ | $e^{-2}$ | $e^{-2}$ | $3.3e^{-2}$ |
| transformer_act_function | GeLU | GeLU | ReLU | ReLU |
| datasets | [filtered Tahoe-100M,<br>filtered CELLxGENE,<br>filtered scBaseCount] | [Tahoe-100M,<br>CELLxGENE,<br>scBaseCount] | [Tahoe-100M,<br>CELLxGENE] | [Tahoe-100M,<br>CELLxGENE] |
| global_batch_size | 10240 | 10240 | 3584 | 400 |
| gradient_accumulation | 40 | 4 | NA | NA |
| #gpus | 32 | 128 | 64 | 16 |
| gpu_type | H200 | H200 | H100 | H100 |

##### Efficiency

The implementation of Tx1 is much more efficient and scalable than similar single-cell foundation model counterparts [[15], [2]], due to the use of the following techniques:

- **StreamingDataset and Streaming DataLoader** We used StreamingDataset and Streaming DataLoader (from the Composer library) instead of PyTorch’s standard DataLoader, which enabled us to reach an MFU as high as ≈ 0.43 for Tx1-3B. Composer’s StreamingDataset is faster mainly because it streams pre-sharded data efficiently from remote storage and fetches it on demand. Instead of downloading entire datasets before training, it streams only the samples needed for the current epoch. This minimizes local I/O bottlenecks and avoids repeatedly preprocessing large datasets. Its optimized I/O pipeline keeps data flowing continuously, unlike the regular DataLoader, which often stalls while waiting for data.
- **Mixed Precision Training** Using mixed precision accelerates training and saves memory because it uses lower-precision (e.g., FP16 or BF16) arithmetic instead of full precision (FP32), which runs much faster on GPUs and reduces memory bandwidth usage.
- **Low-Precision LayerNorm** Low-precision LayerNorm speeds up training by performing normalization operations in reduced precision, lowering compute and memory costs for one of the most frequent operations in transformer blocks.
- **Flash Attention** We introduced a method to train Tx1 without needing the custom masking strategy used in similar methods such as scGPT (explained in Sec. 4.1.3). This strategy enabled us to use FlashAttention for the entire sequence. FlashAttention is faster because it computes attention in chunks directly in GPU SRAM instead of repeatedly reading and writing large attention matrices to slower GPU memory. This drastically reduces memory access and bandwidth costs, making attention computation faster and more memory-efficient.
- **Fully Sharded Data Parallelism (FSDP)** FSDP is faster because it splits model parameters, gradients, and optimizer states across GPUs, so each device only loads the portion it needs at a time. This lowers memory usage and allows us to train the Tx1 series in parallel with less communication overhead.

### 4.3 Evaluation

We evaluated Tx1 series on multiple tasks and benchmarks explained in the following sections.

#### 4.3.1 DepMap Essentiality Prediction

We developed two tasks based on data from the Dependency Map project to evaluate the ability of single-cell foundation models to predict gene essentiality. To set up these tasks, we combined bulk RNA-Seq data from the Cancer Cell Line Encyclopedia (CCLE, [11]) with essentiality data from the Dependency Map (DepMap) project. All raw data was downloaded from the DepMap Portal^5^ as specified below. We considered only the cell lines and genes that were common between the two datasets. Cell lines were matched on DepMap IDs (which have the form ACH-XXXXXX) and genes were matched by official symbol.

- RNA-Seq (fileset: CCLE 2019) `CCLE_RNAseq_genes_counts_20180929.gct.gz` (counts) `Cell_lines_annotations_20181226.txt` (metadata)
- Essentiality (fileset: DepMap Public, version: DepMap Public 23Q4) `CRISPRGeneDependency.csv` (essentiality scores) `Model_v2.csv` (metadata)

##### Predicting broadly essential genes

The goal of this task is to predict whether a gene is broadly essential or inessential, given its embedding. Models are evaluated on held-out genes. We begin by dropping any genes that have NULL essentiality measurements in *>*10% of cell lines and binarizing the remaining essentiality values, where a score ≤ 0.5 is transformed to 0 and is otherwise transformed to 1. We then retain only genes that are essential in *<*5% of cell lines (labeled inessential, *n* = 12,996) or *>*70% of cell lines (labeled essential, *n* = 1,287). These genes are split into five folds for cross-validation, and we train random forest classifiers to predict the inessential / essential binary label from the mean gene embedding across cell lines. Random forests are evaluated using AUROC, which we report across the five folds.

##### Predicting essential genes in each cell line

For this task, the goal is to predict the cell-line-specific essentiality of a gene given the contextual embedding of that gene within a particular cell line. Models are evaluated on held-out cell lines. Since most genes tend to have highly correlated essentiality across cell lines, we systematically split genes across 10%-width strata (e.g., essential in 10–20% of cell lines, essential in 20–30% of cell lines, …) and run models independently for each strata - without this stratification the prediction task would be trivial. Since the 0–10% strata contains broadly inessential genes, we do not use this strata. In the remaining nine strata, we have a median of 159 genes per stratum (minimum 114, maximum 997). Although the task is framed as a regression problem, we binarized both the true and predicted essentiality values and evaluated performance using AUROC rather than *R*^2^. This choice reflects the fact that essentiality scores are typically bimodal - genes tend to be either essential or nonessential. In more detail, we train random forest regressors (using five-fold cross-validation) within each strata, where the input to the model is the contextual embedding of a gene in a cell line, and the target output is the essentiality score for that gene in that particular cell line. We compute the mean AUROC across the five folds in each strata and report the distribution across the strata.

#### 4.3.2 Predicting gene-set membership in MSigDB

We tested the ability of learned gene embeddings to predict gene function by using them to predict gene sets in the molecular signatures database (MSigDB), focusing on the 50 hallmark gene sets [12, 13]. This resulted in a gene-set collection with 4,384 genes. For every gene, we created a multi-hot encoded label corresponding to the gene sets containing the gene. We then trained a multi-layer perceptron with gene embeddings as the input and the signatures as the output using unweighted binary cross-entropy as the loss function.

We extracted the gene embeddings for the MsigDB task for Tx1 series and other models by calculating the mean gene embeddings derived from CCLE expression data (the same dataset used for DepMap essentiality prediction in the previous section). Specifically, each batch of CCLE data is passed through the model, contextual gene embeddings are extracted from the final transformer layer, and then mean-pooled across all available contexts for each gene. Only for TranscriptFormer, the input genes to the model is normalized to 10,000 counts per cell line. And for PCA instead of CCLE dataset we used 500k subset of CZ CELLxGENE.

All resulting embeddings are then passed to the fine-tuning pipeline where we first split the genes into training, validation, and test sets, perform a grid hyperparameter search and early stopping of the MLP on the validation set, and report performance on the test set. We repeated the procedure with five repetitions and reported the performance distribution in Sec. 2.2.2.

### 4.4 Cell-type classification on Tabula Sapiens 2.0

Cell-type classification is a standard benchmark for evaluating how well cell-state embedding models capture transcriptional features associated with cellular identity. We evaluated Tx1 and baseline models using the *czi-benchmarks* package, which provides a standardized leaderboard and evaluation framework for this benchmark.

Because Tx1 and SE-600M were not previously included on the leaderboard, we generated their embeddings and ran them through the benchmark pipeline. This framework computes performance metrics on five tissue groups from the Tabula Sapiens 2.0 dataset - blood, bone marrow, thymus, lung, and mammary gland. The embeddings are used to train three types of classifiers - logistic regression, k-nearest neighbors, and random forest - to predict cell-type annotations. Each classifier is evaluated using five-fold cross-validation, with predictions on held-out data compared against human-annotated cell-type labels to compute accuracy and F1-score. Final scores are averaged across classifiers, folds, and five random seeds to yield a robust measure of model performance. For comparison, results for Transcriptformer and Geneformer-v2-316M are taken directly from the public leaderboard.

### 4.5 State Transition perturbation modeling

A subset of 2,000 previously identified highly variable genes (HVGs) for the Tahoe-100M dataset were extracted to a dense gene expression dataset^6^. The HVG expression was scaled by library size and multiplied by a scaling factor of 1,872 (median library size). In parallel, embeddings for the cells were generated using Tx1 and State Transition. Data were stored as one H5AD file [30] per plate. These data were used to train State Transition models.

Two different data splits for State Transition training: a few-shot split^7^ and a zero-shot split^8^. For each split, three State Transition models were trained using different cell representation spaces: HVG space, State Embedding space, and Tx1 space, using the same number of training steps and other model settings^9^.

#### 4.5.1 State Transition adaptation training

A subset of 2,000 previously identified HVGs for the Parse-PBMC dataset were extracted to a dense gene expression dataset^10^. The HVG expression was scaled by library size and multiplied by a scaling factor of 1,872, then transformed through log_10_(· + 1). In parallel, embeddings for the cells were generated using Tx1 and State Transition.

For adaptation training, the State Transition models were initialized from checkpoints trained on Tahoe-100M for corresponding embedding. Adaptation training was performed for one few-shot split^11^ and one zero-shot split^12^. For each split, three State Transition models were trained using different cell representation spaces: HVG space, State Embedding space, and Tx1 space, using the same number of training steps and other model settings^13^.

#### 4.5.2 Evaluation

Trained models were evaluated on held-out test data using Pearson Δ correlation. Expression deltas (Δ) are defined by the difference in mean expression of a treatment population, 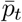, and mean expression of a control population, 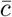, resulting in 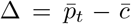. Expression Δ’s are calculated for real (observed) data 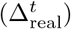 and for predicted data 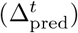. Comparison between the observed and predicted results are quantified by Pearson correlation,

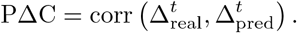

For each dataset and split, predictive performance was quantified with multiple simple baselines representing averaging strategies.

### 4.6 Mean delta baselines

Performance of baseline linear averaging predictions were quantified using different variations of averaging logic, making use of different amounts of experimental design information. Cells can be grouped by context *c*, whether they are from the control condition, or from a perturbation *p*. With this information, per-group averages (or, geometrically, centroids in HVG space) can be defined by

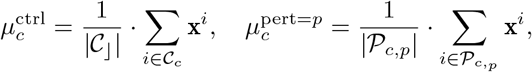

where 𝒞_*c*_ is the set of control cells for context *c* and 𝒫_*c,p*_ is the set of cells with perturbation *p* in context *c*. These averages can be used to produce per-group mean deltas,

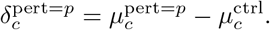

These deltas can be further averaged within the training set of combinations of contexts and perturbations and used for prediction of deltas on the test set, evaluated using PΔC.

Expression delta’s can similarly be calculated separately for individual groupings of cells, such as plates, or other batches. This ensures matching perturbations and controls are appropriately being compared, avoiding spurious correlations due to batch membership. This is achieved by specifying the subset population of cells per group,

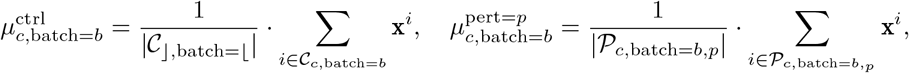

from which we can calculate

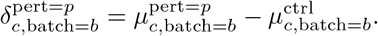

To keep notation simpler, the baselines below are described without grouping, but have also been calculated with grouping.

#### 4.6.1 Null delta baseline

A special case mean baseline assumes 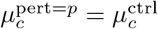, or in other words enforcing *δ* = 0. In this case

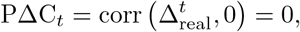

indicating that PΔC_*t*_ is a relative metric, lower-bounded by 0 for predictions with no information.

#### 4.6.2 Global mean delta baseline

The predictive performance of a global mean delta on held out test contexts and perturbations, which has no specific information about either context or perturbation identity, indicates how much systemic differences between the control samples and the remaining samples contributes. The pergroup deltas can be globally averaged to obtain the global mean shift between any treatment and control for any context,

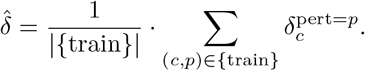

With the global mean delta 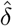, predictions can be made through 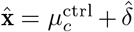 for *c* in the test set. In practice, 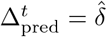 is used to evaluate PΔC_*t*_ on the test set without performing the predictions,

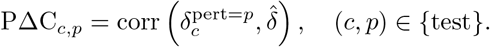

#### 4.6.3 Perturbation mean delta baseline

By creating a mean delta predictor of perturbed expression as control mean for a context shifted by a perturbation-specific offset we make a linear prediction method treating contexts as replicates. Per-perturbation mean deltas can be calculated from the per-group deltas by

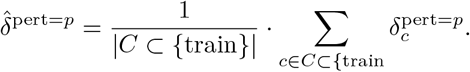

Prediction per context in the test set can be performed by 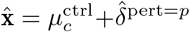. In practice, performance is quantified by

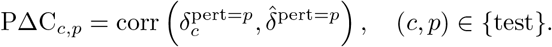

#### 4.6.4 Context mean delta baseline

Conversely, we can predict a general shift of a given cell context to an arbitrary perturbation. These context mean delta shifts will be a linear prediction that treat different perturbations as replicates,

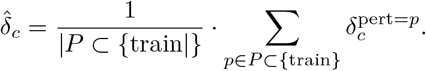

High performance on context mean means that observed responses are uninformed by the specificity of the perturbation. The performance can be computed directly from the averaged averages as

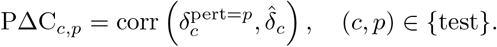

#### 4.6.5 PΔC noise ceiling

The PΔC performance score is based on averaging predicted populations of cells and averaging observed populations of cells, from a single prediction. Segmenting the populations of the observed cells into two disjoint subsets, letting one serve as prediction for the other, will produce a best possible prediction. This way an estimate of the upper limit of possible PΔC performance can be made,

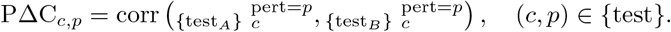

We use the presubscripts {test_*A*_} and {test_*B*_} to indicate that deltas were computed from different subsets of observed test data.

### 4.7 Perturbation separability

Different representations of single cells from single cell foundation models have different ability in reflecting change in transcriptomic change upon perturbation. To quantify this property we define ‘perturbation separability’ as the accuracy of a kNN classifier predicting whether a cell originated from untreated control or treatment, per cell context and treatment.

Before creating the kNN classifier, the number of points is sampled to the minimum of the number of points from control and from treatment. Since the total number of points *N* available varies per context, treatment, and dataset, the number of neighbors *k* is determined dynamically using the rule

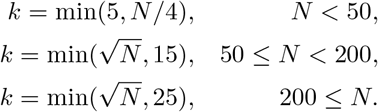

In parallel to calculating the accuracy for treatment and control labels, accuracy is also calculated with a permutation of the labels as a null baseline.

## 5 Data and Code Availability

Pretrained models, fine-tuning scripts, and evaluation datasets are available at https://github.com/tahoebio/tahoe-x1 and https://huggingface.co/tahoebio/TahoeX1

## 6 Acknowledgments

We would like to thank Maya Peters Kostman for their help in designing illustrations for our figures.

## Footnotes

1 https://chanzuckerberg.github.io/cz-benchmarks

2 https://www.parsebiosciences.com/datasets/10-million-human-pbmcs-in-a-single-experiment/

3 https://docs.mosaicml.com/projects/composer

4 https://pytorch.org

5 https://depmap.org/portal/data_page/?tab=allData

6 https://huggingface.co/arcinstitute/ST-Tahoe/blob/main/data_module.torch

7 https://github.com/tahoebio/tahoe-x1/blob/main/scripts/state%20transition/tahoe_5_holdout/generalization_converted_cell_lines_3b.toml

8 https://github.com/tahoebio/tahoe-x1/blob/main/scripts/state%20transition/tahoe_5_holdout/zeroshot_generalization_converted_cell_lines_3b.toml

9 https://github.com/tahoebio/tahoe-x1/tree/main/scripts/state%20transition

10 https://huggingface.co/arcinstitute/ST-Parse/blob/main/data_module.torch

11 https://github.com/tahoebio/tahoe-x1/blob/main/scripts/state%20transition/parse_pbmc_holdout/donor_tx13b_parse_hvg_v3.toml

12 https://github.com/tahoebio/tahoe-x1/blob/main/scripts/state%20transition/parse_pbmc_holdout/donor_zeroshot_tx13b_parse_hvg_v3_embs.toml

13 https://github.com/tahoebio/tahoe-x1/tree/main/scripts/state%20transition

